# Personal care formulations demonstrate virucidal efficacy against multiple SARS-CoV-2 variants of concern: implications for hand hygiene

**DOI:** 10.1101/2021.09.20.461073

**Authors:** Sayandip Mukherjee, Carol K. Vincent, Harshinie W. Jayasekera, Ashish Shrikant Yekhe

## Abstract

The second and third waves of COVID-19 pandemic have largely been driven by the surge of successive SARS-CoV-2 variants of concern (VOC). These VOC have rapidly spread through multiple geographies being enabled by high transmission rates and/or high viral load compared to the original parent strain. Consequently, the altered phenotypes of these VOC have posed greater challenges to diagnostic and clinical management of COVID-19. Despite considerable progress being made on vaccine roll out, practicing proper hand hygiene has been advocated as a consistent precautionary intervention as more virulent VOC continue to emerge and spread across geographies.

Two variants of concern, namely beta and delta, have recently been shown to escape antibody-mediated neutralization by virtue of acquired mutations in the receptor-binding domain of the viral spike protein which binds to the human ACE2 receptor for cellular entry. In this report we have empirically determined the efficacy of a range of personal care formulations in inactivating the beta and delta variants of SARS-CoV-2. High titres of these variants were exposed to marketed personal care formulations from Unilever under standard in-vitro suspension test-based conditions relevant to end-user habits. All the formulations demonstrated greater than 99.9% reduction in viral infective titres. The rate of inactivation by these products were comparable to that of the original strain of SARS-CoV-2 virus tested under the same conditions. Therefore, it can be concluded that well-designed personal care formulations when tested under consumer-centric conditions, and with proven efficacy against the parent strain of SARS-CoV-2 will continue to be effective against extant and emerging variants of SARS-CoV-2. This is through their broad-spectrum mode of action (disruption of lipid bilayer of the host-derived viral envelope, denaturation of envelop and nucleocapsid proteins, and disruption of genome) which is independent of the escape mutations that facilitate immune evasion or enhanced transmissibility.

## Introduction

Generation of successive variants is an expected consequence of viral evolution and results primarily from extended viral persistence and propagation in the human population. While majority of genetic variants are of insignificant consequence to the human host, certain traits of a selected number of variants could prolong the pandemic and reintroduce risk of COVID-19 even among vaccinated individuals [1, 2]. The World Health Organization (WHO) defines classes of SARS-CoV-2 variants based on severity of illness, transmissibility, and their impact on public health and social countermeasures such as lockdowns and masking, and medical countermeasures such as diagnostics, therapeutics, and vaccination. Variants of Concern (VOC) are those with evidence of inducing more severe disease, increasing transmissibility, or having negative impact on countermeasures. There are currently four VOC designated by the WHO as Alpha, Beta, Gamma, and Delta (https://www.who.int/en/activities/tracking-SARS-CoV-2-variants/). In an analysis of the global spread of SARS-CoV-2 variants and estimated changes in the effective reproduction numbers (R_0_) at country-specific level, it was clearly demonstrated that WHO-designated VOC have rapidly replaced previously circulating lineages with estimated transmissibility increases of 29%, 25%, 38% and 97% respectively for Alpha, Beta, Gamma, and Delta [3]. In addition to enhanced transmissibility, many of these VOC have shown increased potential for immune evasion / escape. Multiple VOC, including those such as the Beta variant with substitutions at residues 484 (E484K) and 417 (K417N), have demonstrated escape to neutralization by monoclonal antibodies directed to the ACE2-binding Class 1 and the adjacent Class 2 epitopes but are susceptible to neutralization by the generally less potent antibodies directed to Class 3 and 4 epitopes on the flanks of the receptor binding domain (RBD) [4] [5]. Reinfection by Beta variant is also possible after COVID-19 as neutralization by convalescent serum was found to be significantly reduced [6]. Similarly, in-depth studies with the Delta variant have shown that reduction in neutralization titres using convalescent or vaccine sera may lead to some breakthrough infections. Additionally, the study results suggest that a proportion of unvaccinated individuals infected previously with Beta and Gamma variants may be at higher risk of reinfection with Delta [7].

Although vaccination remains the main-stay of protection from COVID-19, and extensive research is currently underway to understand the correlates of protection against SARS-CoV-2 and the circulating variants, health authorities have continued to emphasize the role of proper hand and respiratory hygiene in reducing the risks of infection and transmission among the susceptible population [8, 9]. Proper hand-hygiene is considered one of the most accessible and economic public health measures to reduce transmission of germs across community settings. Multiple systematic reviews and meta-analysis have firmly established the role of good hand hygiene in low-, middle-, and high-income countries in reducing incidence of acute respiratory tract infections including seasonal influenza and viral pneumonia [10-12]. In similar lines we have previously established that marketed personal care formulations such as bar soap, liquid handwash, and alcohol-based sanitizer from Unilever are capable of greater than 99.9% reduction of infective viral titre of the primary SARS-CoV-2 strain when tested under in-vitro conditions mimicking real life usage conditions [13]. In this report, we seek to establish and re-affirm the efficacy of selected and previously tested personal care formulations in inactivating two major VOC, namely Beta and Delta, under standardized test methods with conditions relevant to end-user.

## Materials and Method

### Viruses and cell culture

Severe Acute Respiratory Syndrome Coronavirus 2 (SARS-CoV-2) strains hCoV-19/South Africa/KRISP-EC-K005321/2020 (PANGO lineage B.1.351 / Beta VOC) and hCoV-19/USA/PHC658/2021 (PANGO lineage B.1.617.2 / Delta VOC) were sourced from BEI Resources. Vero-E6 cells were obtained from ATCC (#CRL-1586).

### Test products

The following commercially available personal care formulations from Unilever were tested – proprietary soap bar with minimum 30% (w/w) total soluble surfactant (soluble soap with added soluble surfactant); proprietary liquid cleansers (handwashes) with surfactant concentrations of 7.8% (w/w) and 12% (w/w); proprietary alcohol-based sanitizer containing 65% alcohol (w/w).

### Assay for virus Inactivation

Fully formulated products were evaluated in suspension for virucidal efficacy based on the ASTM International Standard E1052-20. Bar cleansers were prepared as 8% w/v solutions and assessed at 40° + 2°C for 20 seconds contact time. Liquid cleansers were prepared as 50% w/v solutions while hand sanitizers were evaluated neat; both were tested at 20° + 2°C for 20- and 10-seconds contact time respectively. Each test product was tested in triplicate. All controls (virus recovery, neutralization, and cytotoxicity) were performed according to ASTM standard. Host cells were examined microscopically for virus-specific cytopathic effects and product-specific cytotoxic effects.

### Measurement of titre

The 50% tissue culture infective dose per mL (TCID_50_/mL) was determined using the Spearman-Karber method, and in cases where no virus was detectable, a statistical analysis using the Poisson distribution was performed to determine the theoretical maximum possible titre for the product. The average log_10_ reduction was calculated as the difference in TCID_50_ between the test product and virus recovery control.

## Results

### Inactivation of SARS-CoV-2 variants by personal care products

The virucidal efficacy of three classes of fully formulated commercial products namely bar soap, liquid handwash, and alcohol-based sanitizer was tested against two SARS-CoV-2 VOC namely Beta and Delta. As shown in table 1, soap bar gave greater than equal to log_10_ 3.4 (≥ 99.9%) reduction of SARS-Cov-2 Beta variant titre and greater than equal to log_10_ 3.2 (≥ 99.9%) reduction of SARS-Cov-2 Delta variant titre. Next, we investigated the efficacy of liquid handwash with two different total surfactant contents of 7.8% and 12% and found that they reduced the input titre of Beta variant by greater than equal to log_10_ 3.7 (≥ 99.9%) and by greater than equal to log_10_ 3.1 (≥ 99.9%) of Delta variant. The contact duration of both the above sets of experiments were in accordance with the WHO’s recommendation of 20 seconds of handwashing.

**Table 1.** Virucidal efficacy of a range of commercially available proprietary personal care formulations tested against selected SARS-CoV-2 variants of concern (VOC) at user relevant dilutions and contact duration. TCID_50_, 50% tissue culture infective dose.

| SARS-CoV-2 VOC | Product format | Product Description | Test temperature | Test concentration | Contact time | Initial viral load (Log<sub>10</sub> TCID<sub>50</sub> / mL) | Output viral load (Log<sub>10</sub> TCID<sub>50</sub> / mL) | Log<sub>10</sub> reduction |
| --- | --- | --- | --- | --- | --- | --- | --- | --- |
| Beta (PANGO lineage B.1.351) | Bar soap | Minimum 30% (w/w) total soluble surfactant | 40° ± 2°C | 8% | 20 sec | 7.56 | ≤ 4.13 | ≥ 3.43 |
|  | Liquid cleanser | 7.8% surfactants (w/w) | 20° ± 2°C | 50% | 20 sec | 7.37 | ≤ 3.61 | ≥ 3.76 |
|  | Alcohol based Sanitizer | 65% alcohol (w/w) | 20° ± 2°C | 100% | 10 sec | 7.04 | ≤ 2.61 | ≥ 4.43 |
| Delta (PANGO lineage B.1.617.2) | Bar soap | Minimum 30% (w/w) total soluble surfactant | 40° ± 2°C | 8% | 20 sec | 7.39 | ≤ 4.13 | ≥ 3.26 |
|  | Liquid cleanser | 12% surfactants (w/w) | 20° ± 2°C | 50% | 20 sec | 7.71 | ≤ 4.61 | ≥ 3.10 |
|  | Alcohol based Sanitizer | 65% alcohol (w/w) | 20° ± 2°C | 100% | 10 sec | 5.87 | ≤ 2.61 | ≥ 3.26 |

In the absence of soap and water, the CDC recommends the use of alcohol-based hand rub containing at least 60% alcohol (w/w) for hand disinfection. Therefore, we evaluated alcohol-based hand sanitizer containing 65% alcohol (w/w) for virucidal efficacy against the SARS-CoV-2 VOC. As noted previously with hand washing, we believe that even for alcohol-based hand sanitizers it is critical to look at lower contact time-points which are more consumer relevant. In an observation of 50 episodes of hand hygiene with an alcohol-based hand rub of health care personnel working in an intensive care unit, it was recorded that the mean time of application (i.e., rubbing product onto hands) was 11.6 seconds with a median time of 10 seconds[14]. As shown in table 1, the 65 % alcohol-based sanitizer gave greater than equal to log_10_ 4.4 (≥ 99.99%) reduction of SARS-CoV-2 Beta titre in 10 seconds while the same sanitizer achieved greater than equal to log_10_ 3.2 (≥ 99.9%) reduction for Delta over the same contact duration. It is worthwhile to mention here that this discrepancy can be accounted for by the lower input titre that was employed for the Delta variant. All the test products reduced the infective titre of SARS-CoV-2 VOC below detectable limits by greater than 99.9% within 10 to 20 seconds of exposure and at dilutions relevant to end-user usage.

## Discussion

As of September 6, 2021, reported cases of infection by Delta variant have crossed the million-mark having spread across 150 countries and accounting for almost 100% of new infections detected in the UK, USA and Denmark (data from GISAID). Here we established the virucidal efficacy of a range of marketed personal care products from Unilever against both the Beta and Delta variants of SARS-CoV-2 and found them to be comparable to previously demonstrated efficacy of similar range of personal care products against the original SARS-CoV-2 strain. To the best of our knowledge, this is the first study to report inactivation of Beta and Delta VOC by commercially formulated personal care products for cleansing. Our results re-affirm the importance of hand-hygiene products with adequate levels and composition of surfactants (for soap bars and liquid cleansers) and alcohol (for sanitizers) to deliver efficacy in consumer habit-centric conditions to prevent further spread of VOC. We believe that these results will lend confidence to members of the public and encourage them to continue to practice good hand hygiene. However, we should not lose sight of the fact that the world at this point is closely monitoring the spread and behaviour of multiple variants of interest (VOI) namely Eta, Iota, Kappa, Lambda and Mu, and numerous variants of high consequence, all of which have the potential to become future VOC[15]. Therefore, it is important to consider whether daily hygiene measures involving handwashing and hand-sanitation with soaps and alcohol-based hand-rubs should be expected to be effective against emerging VOC. To answer this, we must look at the mechanism of action as related to viral mutations and their effects on fundamental virus particle structure and function.

Coronaviruses including SARS-CoV-2 are multimolecular assemblies of virally derived nucleic acids (RNA), proteins (spike, membrane, envelope and nucleocapsid), and host derived lipids (viral envelope). The nucleocapsid which encloses the viral genomic RNA is surrounded by the lipid bilayer in which the spike proteins, membrane glycoproteins, and envelope proteins are anchored. The different constituents of a mature virion are held together by non-covalent interactions. These non-covalent interactions are weak in nature and the virus relies on the host-derived envelope for its shape, integrity, and infectivity.

Surfactants present in soaps, liquid handwashes and detergents, are amphipathic molecules with a hydrophobic fatty acid tail and a hydrophilic head group. When the surfactants encounter the virus, they insert themselves into the lipid bilayer aided by the hydrophobic tails while the hydrophilic head groups remain outside in contact with water. The key tendency of a micelle-forming amphiphile inserting into a lipid bilayer is its preference for a locally curved interface (its spontaneous curvature) that conflicts with the planar topology of a bilayer. This misfit causes a curvature stress. As more and more surfactant molecules wedge themselves into the ordered lipid bilayer, the viral envelope is ruptured thereby destroying the multimolecular assembly and rendering the virus inactive. When the surfactant concentration reaches the critical micellar concentration (CMC), micelles form around the disintegrated viral particles and remove them during rinsing[16, 17].

Most alcohol-based sanitizers contain either ethanol, 1-propanol, 2-propanol (IPA), or a combination of these molecules dissolved in water. Alcohol-based products inactivate SARS-CoV-2 virus particles by disrupting the structure of the proteins (a process called denaturing) on the surface of the virus. On exposure, alcohols displace the hydrogen bonds between amino acids that hold the viral proteins in shape, causing the proteins to lose their structure and function, thereby inactivating the virus. The water present in majority of hand sanitizers including the one studied here, plays a key role as proteins are difficult to disrupt by this method in the absence of water. This means that alcohol solutions are most effective when they contain 60–80% alcohol rather than 100%. Alcohols also disrupt the virus lipid membrane, but this occurs by a different mechanism. The higher the lipid content and the larger the virus particle, the more susceptible the virus is to alcohols. The alcohols used in sanitizers are small polar molecules that can interact with the surface of the lipid membrane. When alcohols are present in sufficient concentrations, they disrupt the ordered structure of the membrane, breaking open the viral particle[18-20].

Many other broad-spectrum virucidal agents such as organic acids, quaternary ammonium compounds, phenolics, and oxidizers are also known to disrupt the viral envelope, the viral genome, and the viral envelope proteins through physico-chemical modes of action. The representative everyday hand hygiene products studied here do not contain such agents, but it would be expected that effective formulations containing the above-mentioned agents would also be minimally impacted by viral mutations given similar mechanisms of action to those ingredients discussed here.

Majority of the lineage-defining mutations occur in the spike protein for both VOC and VOI. These mutations are chiefly single-nucleotide substitutions which alter the affinity of the spike protein for the ACE2 receptor and/or for neutralizing antibodies. While these mutations affect binding of the RBD to the receptor or antibodies at the molecular level, they do not alter the macromolecular property of spike protein which therefore remains susceptible to denaturation by soaps, surfactants, alcohols, organic acids, phenolic compounds and oxidizers. Similarly, the lipid components of the host-derived viral envelope remain completely permissive to disruption by surfactants and alcohol as they are unaffected by the mutations and remain unchanged across the variants.

In conclusion, we have provided empirical proof that marketed personal care formulations from Unilever have high virucidal efficacy against the SARS-CoV-2 variants Beta and Delta. This direct experimental evidence fully supports the broader argument that well-designed personal care hand hygiene products capable of delivering efficacy in user centric conditions which act through physico-chemical mechanism against the basic structure of the virus particle should be efficacious regardless of mutations [21]. Furthermore, we argue that due to broad-spectrum mode of action of these tested formulations, the continued practice of good hand hygiene practices with everyday used products such as those described herein holds significant promise as an easily accessible, economic and effective non-therapeutic intervention towards reducing the transmission of present and future variants of SARS-CoV-2 across communities and populations.

## Acknowledgements

We would like to thank Microbac Laboratories, Sterling, Virginia, USA for providing technical assistance. The authors would also like to thank Dr. Timothy Tobery and Dr. Naresh Ghatlia for critical review of the manuscript.

